# 14-3-3 Negatively Regulates Actin Filament Formation in the Deep Branching Eukaryote *Giardia lamblia*

**DOI:** 10.1101/142505

**Authors:** Jana Krtková, Jennifer Xu, Marco Lalle, Melissa Steele-Ogus, Germain C. M. Alas, David Sept, Alexander R. Paredez

## Abstract

The phosphoserine/phosphothreonine-binding protein 14-3-3 is known to regulate actin, this function has been previously attributed to sequestration of phosphorylated cofilin. The deep branching eukaryote *Giardia lamblia* lacks cofilin and all other canonical actin-binding proteins (ABPs), and 14-3-3 was identified as an actin-associated protein in *Giardia*, yet its role in actin regulation was unknown. Gl14-3-3 depletion resulted in an overall disruption of actin organization characterized by ectopically distributed short actin filaments. Using phosphatase and kinase inhibitors, we demonstrated that actin phosphorylation correlated with destabilization of the actin network and increased complex formation with 14-3-3, while blocking actin phosphorylation stabilized actin filaments and attenuated complex formation. *Giardia's* sole Rho family GTPase, GlRac, modulates Gl14-3-3's association with actin, providing the first connection between GlRac and the actin cytoskeleton in *Giardia*. *Giardia* actin contains two putative 14-3-3 binding motifs, one of which (S330) is conserved in mammalian actin. Mutation of these sites reduced, but did not completely disrupt, the association with 14-3-3. Native gels and overlay assays indicate that intermediate proteins are required to support complex formation between 14-3-3 and actin. Overall, our results support a role for 14-3-3 as a negative regulator of actin filament formation.

**Importance:** *Giardia* lacks canonical actin binding proteins. 14-3-3 was identified as an actin interactor but the significance of this interaction was unknown. Loss of 14-3-3 results in ectopic short actin filaments, indicating that 14-3-3 is an important regulator of the actin cytoskeleton in *Giardia*. Drug studies indicate that 14-3-3 complex formation is in part phospho-regulated. We demonstrate that complex formation is downstream of Giardia’s sole Rho family GTPase, GlRac, this result provides the first mechanistic connection between GlRac and actin in *Giardia*. Native gels and overlay assays indicate intermediate proteins are required to support the interaction between 14-3-3 and actin suggesting that 14-3-3 is regulating multiple actin complexes. Overall, we find that 14-3-3 is a negative regulator of actin filament formation in *Giardia*.

## Introduction

14-3-3 belongs to a family of highly conserved eukaryotic proteins whose role is to regulate target proteins through binding of specific phosphoserine/phosphothreonine motifs. Through recognition and binding of these specific motifs, 14-3-3 functions in a variety of cellular processes, including cytoskeletal regulation and function as an adapter protein that can activate/inhibit protein function, change intracellular localization of bound cargos, or mediate formation of multi-protein complexes (1-8). In the current model for higher eukaryotes, 14-3-3 regulates actin through phospho-dependent sequestration of the actin-depolymerizing protein, cofilin (4, 9). The existence of multiple 14-3-3 isoforms in higher eukaryotes complicates the relationship between 14-3-3 and actin, leading to discrepant results about whether 14-3-3 directly interacts with actin (2). Consistent with additional interaction/regulatory mechanisms, actin has been identified as an interactor of 14-3-3 in plant and animal 14-3-3 proteomic datasets where cofilin was not found (10-12). Indeed, 14-3-3σ was recently reported to be upregulated in breast cancer cells, where it forms a complex with actin and intermediate filament proteins that is utilized for cell motility during breast tumor invasion (13). The complex was also found to play a role in actin sequestration, as depletion of 14-3-3σ led to an increase in filamentous actin. Whether complex formation between monomeric actin and 14-3-3 is broadly utilized mechanism of actin regulation remains an open question.

*Giardia lamblia* (synonymous with *G. intestinalis* and *G. duodenalis)*, is a protozoan parasite that belongs to a deep-branching group of eukaryotes known as Excavata. *Giardia*, in concordance with its phylogenetic position, has an evolutionarily divergent actin with only 58% average identity to other actin homologs and lacks the canonical actin-binding proteins once thought common to all eukaryotes (Arp2/3 complex, formin, wave, myosin, cofilin, etc.) (14-16). Other excavates, such as *Trichomonas vaginalis* and *Spironucleus salmonicida*, also lack many canonical actin-binding proteins suggesting that the core actin regulators conserved in plants, animals, and fungi may not have solidified their cellular roles before the ancestors of these excavates branched from the eukaryotic tree (17-20). Nevertheless, *Giardia* actin (GlActin) functions in conserved cellular processes including membrane trafficking, cytokinesis, polarity, and control of cellular morphology (21). The mechanism for actin recruitment and regulation for these processes remains poorly understood. The only conserved actin regulator identified in *Giardia* is a Rho family GTPase, GlRac, which can promote changes in actin organization without any of the actin-binding proteins known to link small G-protein signaling to the actin cytoskeleton (21). Notably, 14-3-3 has been shown to integrate G-protein signaling to the actin and tubulin cytoskeleton in *Dictyostelium discoideum* (7); thus, it potentially links GlRac to the actin cytoskeleton in *Giardia*. Through actin affinity chromatography and MudPIT analysis, the single 14-3-3 homolog (Gl14-3-3) of *Giardia* was identified as an actin-associated protein (19). Likewise, actin has been identified as part of the 14-3-3 interactome in *Giardia* (22). Here we set out to address whether 14-3-3 has a role in regulating the actin cytoskeleton, characterize the nature of the interaction, and determine if *Giardia′s* sole Rho family GTPase, GlRac, is upstream of this association.

## Results

We previously reported that 14-3-3 complexes with actin and the biochemical conditions used to isolate interactors suggested the interaction was likely with monomeric G-actin (19). Pulldown of TwinStrep-tagged GlActin (TS-actin) supports this assertion. Cleared lysates (Figure 1A) contain both endogenous GlActin and TS-actin. If 14-3-3 bound to F-actin or complexes containing dimeric actin, then the filaments would contain a mixture of native and tagged actin. After pulldown with StrepTactin resin in buffers not expected to support filament formation, only TS-actin was detected (Figure 1A). This finding is consistent with the 14-3-3 complex containing monomeric actin, but does not exclude interaction with F-actin.

**Figure 1.**
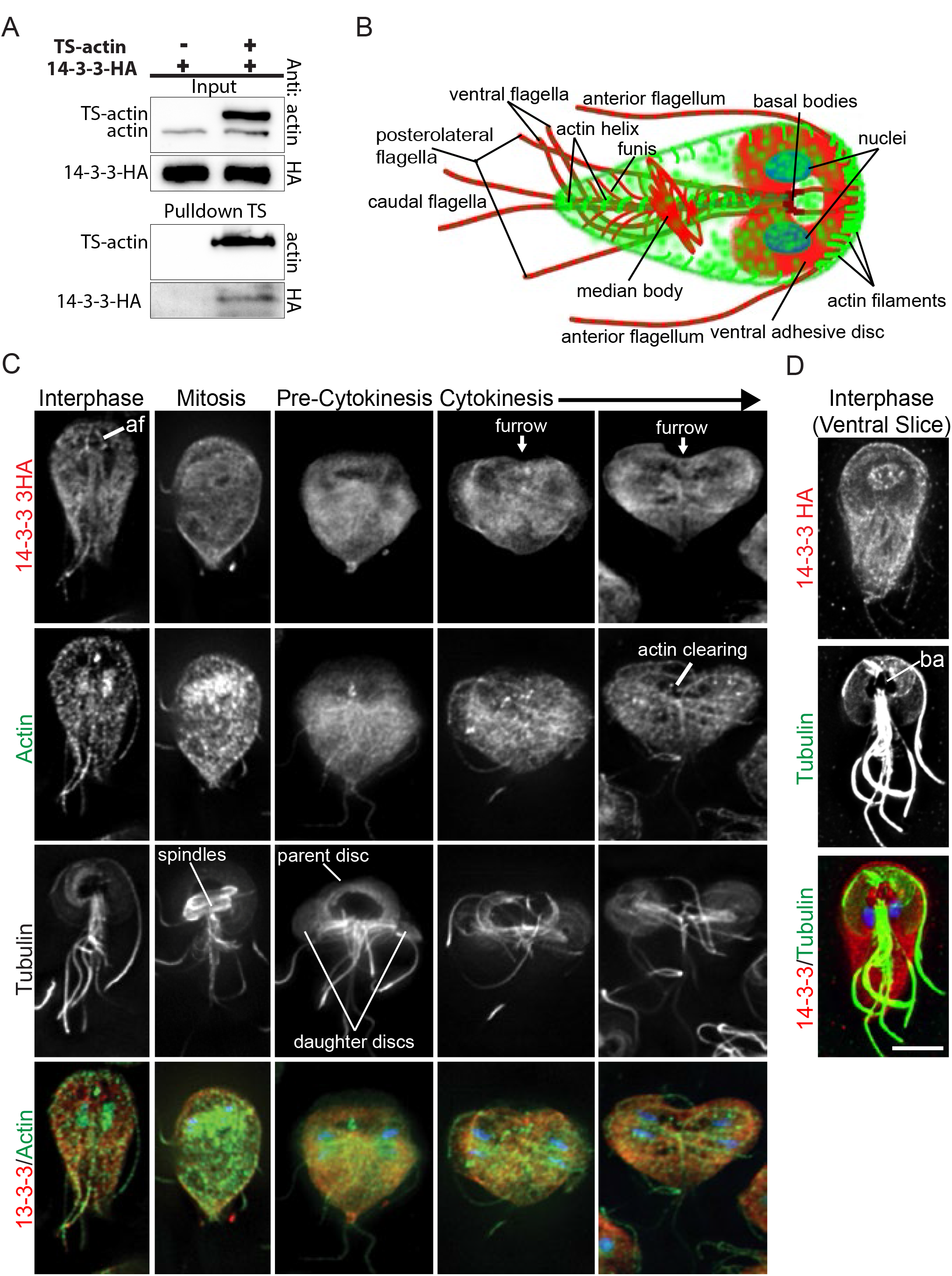
14-3-3 is associated with monomeric actin. (A) Pulldown of TS-actin demonstrating that 14-3-3 interacts with monomeric actin. (B) Diagram of actin (green) and tubulin (red) cytoskeletal structures found in interphase *Giardia* trophozoites. (C) Gl14-3-3-HA (red), GlActin (green), tubulin (greyscale), and DNA (blue) localized in interphase, mitosis and cytokinesis. Gl14-3-3-HA was enriched along the intracytoplasmic portion of the anterior flagella (af). (D) Gl14-3-3-HA (red), tubulin (green), and DNA (blue) projection spanning the ventral region only. Note Gl14-3-3-HA in the microtubule bare area (ba) of the ventral disc (conduit for membrane trafficking). Scale bar= 5μm.

Actin levels in *Giardia* have yet to be measured; assignment of the intracellular concentration relative to the critical concentration has important regulatory implications. If the concentration of actin is above the critical concentration, then *Giardia* would require a mechanism to sequester actin. We questioned whether there would be sufficient 14-3-3 to bind and modulate actin function. Using purified proteins as standards and custom antibodies to GlActin and Gl14-3-3, we measured actin and 14-3-3 concentrations in *Giardia* trophozoite extracts. We found that 10 μg of extract contained 102.5±7.4 ng of Gl14-3-3 and 70.7± 16.4 ng of GlActin or ~1.8 picomoles of 14-3-3 dimer and ~1.7 picomoles of actin (Figure S1). Our measurement of actin at 70 ng per 10 μg of total cellular extract can be extrapolated to ~4.7 μM actin (16,927 cells=10 μg; 1cell=199.8μm^3^ (23)). Compared with other eukaryotes, this actin concentration is relatively low; yet, the value is at least 5x higher than the concentration needed to form filaments (21), indicating that some level of actin sequestration is likely needed to properly regulate filament formation.

Since 14-3-3 has a role in regulating cell division in other eukaryotes, we examined the localization of an endogenously C-terminally tagged version of Gl14-3-3 (Gl14-3-3-HA)(19); see Figure 1B for a diagram of *Giardia* cellular landmarks. In interphase cells, 14-3-3 is distributed throughout the cell with some enrichment at the cortex, nuclear envelope, and in association with the intracytoplasmic axonemes of the anterior flagella. In mitotic cells, 14-3-3 disassociated with the intracytoplasmic axonemes but maintained association with the nuclear envelopes/spindles (Figure 1C). Notably, we previously demonstrated a central role for actin in positioning the flagella and nuclei (21). Gl14-3-3 was also associated with the ingressing furrow, which does not utilize a contractile ring (Figure 1C). We recently reported that actin levels are reduced just ahead of the advancing furrow cortex, and actin is required for abscission but not furrow progression (Hardin et al, in review). Enrichment of Gl14-3-3 just ahead of the furrow cortex may indicate a negative actin regulatory function for Gl14-3-3 and/or a role in regulating membrane trafficking (Figure 1C). Consistent with 14-3-3 having a role in regulating membrane trafficking (6), Gl14-3-3-HA is associated with the nuclear envelope/ER and the bare area of the ventral disc (Figure 1D). This void in the disc lacks microtubules and serves as a conduit for vesicle trafficking whereas the rest of the disc is composed of a sheet of microtubules and associated proteins that would physically prevent vesicle transport (24). The Gl14-3-3-HA fusion protein did not appear to co-localize with filamentous actin (F-actin) structures and is consistent with our finding that Gl14-3-3 complexes with monomeric actin.

To ascertain whether Gl14-3-3 has a role in cytoskeletal regulation in *Giardia*, we depleted Gl14-3-3 with an antisense translation-blocking morpholino. Knockdown of Gl14-3-3 protein expression was monitored by immunoblotting to detect the integrated copy of Gl14-3-3-HA. On average, a 70% reduction in Gl14-3-3-HA levels was observed 24 hours after morpholino treatment and parasite growth was dramatically reduced in the knockdown population versus the non-specific morpholino control, indicating a key role in cell proliferation (Figure 2A, B). Depletion of Gl14-3-3 disrupted characteristic actin organization and resulted in small bright puncta distributed throughout the cell (Figure 2C, S2). Detailed examination revealed that the puncta are short filaments below 1 μm in length (Figure 2D, Movie S1). Depletion of Gl14-3-3 also led to polarity and cytokinesis defects, indicating that Gl14-3-3 has a role in regulating both the actin and tubulin cytoskeletal organization (Figure 2E, S2). The loss of cell polarity, accumulation of multinucleate cells, reduction in cell growth, and abnormal flagella positioning associated with Gl14-3-3 depletion are phenotypes also observed in GlActin depleted *Giardia* (21). These results strongly indicate that 14-3-3 is linked to actin regulation in *Giardia*.

**Figure 2.**
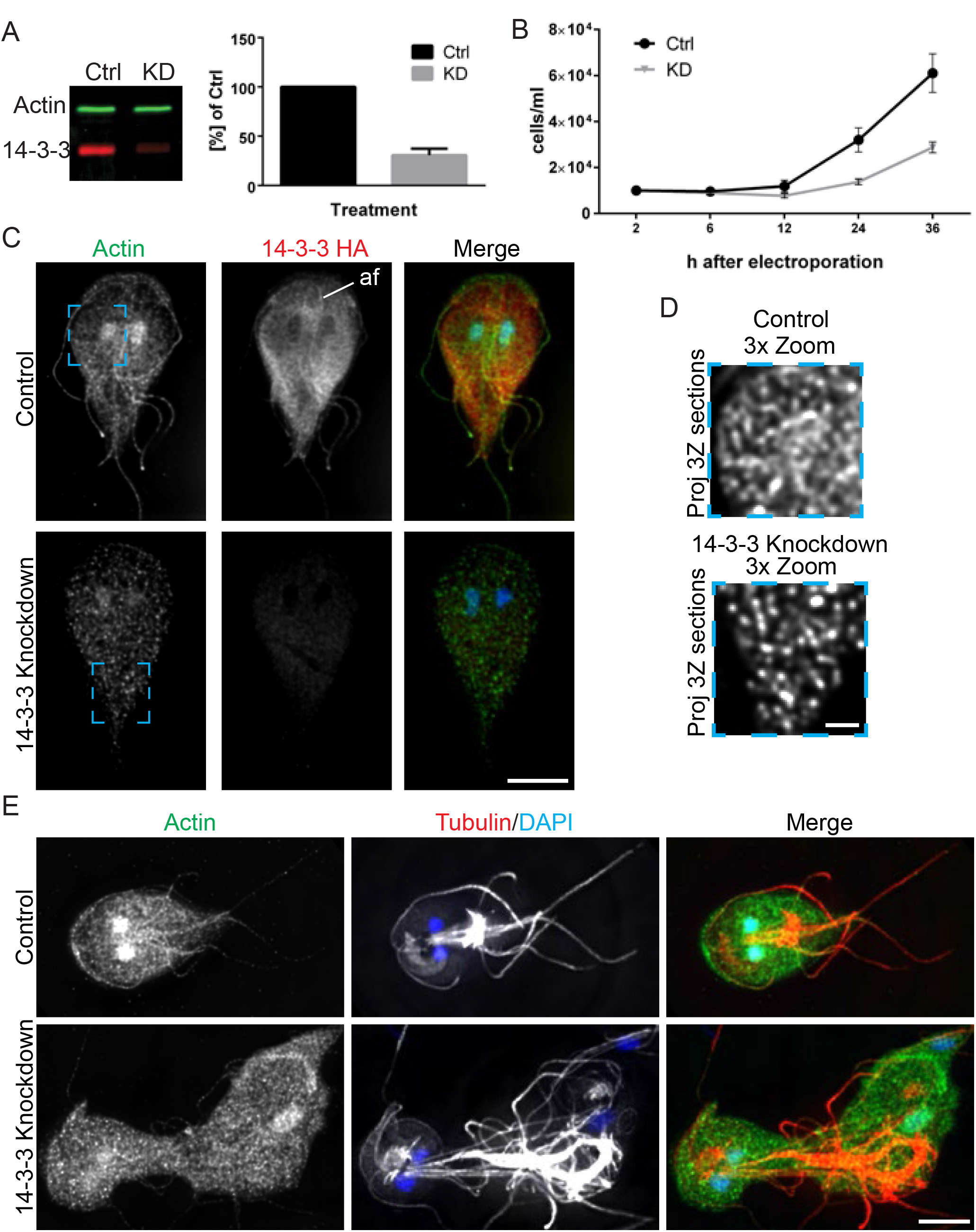
14-3-3 is required for *Giardia* actin cytoskeletal organization and growth. (A)howing typical Gl14-3-3-HA reduction 24 hours after morpholino treatment and quantification of three independent experiments. (B) Growth curves of control and Gl14-3-3 depleted cell cultures indicate that Gl14-3-3 is critical for *Giardia* culture growth (Error=SD). (C) Immunofluorescence staining of control and Gl14-3-3 depleted cells scaled equally. Note enrichment of Gl14-3-3-HA along the intracytoplasmic axonemes of the anterior flagella (af) and that depletion of Gl14-3-3 altered actin organization. Scale bar=5 μm. (D) A magnified view of the blue box in C, optimally scaled to show actin filaments in the control and Gl14-3-3 depleted cells. The puncta in Gl14-3-3 depleted cells are short filaments; see Movie S1 for an entire image stack. Scale bar= 1 μm. (E) Actin (green) and Tubulin (red) staining show 14-3-3 depleted cells lose cell polarity and have cytokinesis defects. See figure S1 for further examples of knockdown phenotypes. Scale bar= 5 μm.

Since actin phosphorylation occurs in several eukaryotes (25-31) and 14-3-3 binding usually requires Ser/Thr target phosphorylation, the effect of kinase and phosphatase inhibitors on actin/14-3-3 complex stability was studied. The Ser/Thr phosphatase inhibitor calyculin A and the general kinase inhibitor staurosporine are both effective in *Giardia* (32), likely affecting the phosphorylation state of multiple proteins. Using Phos-Tag phosphate-affinity electrophoresis (33), we find that a portion of actin is indeed phosphorylated in *Giardia* extracts, (Figure 3A), suggesting phosphorylation could be an important actin regulatory mechanism. After 45 minutes of treatment with either staurosporine or calyculin A, the level of actin phosphorylation increased following treatment with the phosphatase inhibitor and decreased as a result of treatment with the kinase inhibitor, respectively (Figure 3A). Treatment of the cell extracts with lambda protein phosphatase effectively depletes the shifted band demonstrating the specificity of our actin antibody. Remarkably, our ability to co-immunoprecipitate actin with Gl14-3-3-HA correlated with the phosphorylation level of actin and is consistent with phosphor-dependent regulation of the 14-3-3-actin interaction (Figure 3B, 3C).

**Figure 3.**
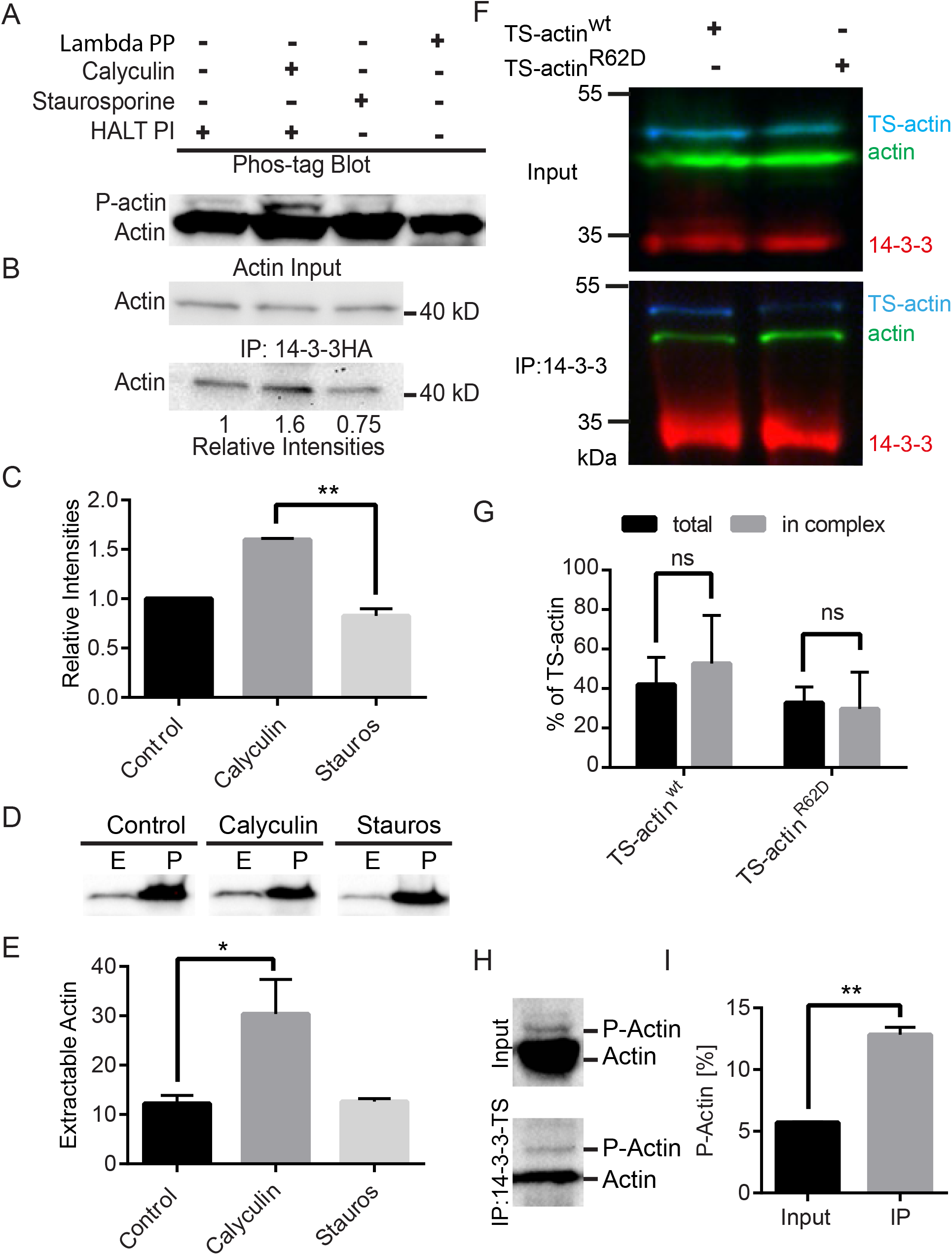
14-3-3-Actin complex formation is phosphodependent. (A) Immunoblot after Phos-tag phosphate-affinity electrophoresis. Cells were pre-treated with DMSO or inhibitors and then HALT phosphatase inhibitor (HALT PI) was added at lysis to preserve the phosphorylation state. Calyculin A treatment increased phosphorylated-actin levels (P-actin) and the kinase inhibitor staurosporine reduced P-actin. Phosphoisoforms were removed after lambda protein phosphatase treatment. (B) Immunoprecipitation of Gl14-3-3-HA after calyculin A treatment led to increased actin interaction while staurosporine treatment reduced the association of actin with Gl14-3-3-HA. (C) Mean values of three independent experiments, error bars=SD and P<0.01. (D) Detergent extractable actin (E=extracted, predominantly G-actin); P=cell pellet/nonextracted, predominantly F-actin) is increased by calyculin A treatment. (E) Plots are mean percentage of extractable actin from three independent experiments, error=SD, P<0.05. (F) Pulldown of 14-3-3 in cells expressing wild type TS-actin or the polymerization defective R62D mutant. (G) Graph showing binding of wild type TS-actin compared to R62D polymerization defective mutant in three independent experiments, ns=not statistically significant. (H) Compared with input, eluted protein from 14-3-3-TS pulldown show enrichment of P-actin. (I) Quantification of three independent experiments, P<0.01.

Next we asked whether modulating the phosphorylation level of actin could affect the balance between F and G-actin. Actin extraction assays were performed after treatment of parasites with staurosporine or calyculin A. Phosphatase inhibition with calyculin A increased extractable actin from 12.2% in DMSO treated control cells to 30.4% (n=3, p<0.05). No reduction in extractable actin was observed after treatment with the kinase inhibitor staurosporine (Figure 3D, 3E), possibly due to the limited sensitivity of this assay coupled with our observation that only a small pool of actin is free to begin with. These results do show that actin phosphorylation is correlated with a shift in the balance toward soluble, presumably, G-actin and association with 14-3-3.

The increased association of Gl14-3-3 with actin that results from calyculin A treatment could indirectly result from increased monomeric actin levels. Therefore, we sought to assess whether increased monomeric actin in the absence of increased phosphorylation could promote complex formation with Gl14-3-3. Mutation of Arg 62, a key residue also conserved in GlActin, to Asp (R62D) has been shown to result in polymerization defective β-actin (34). To monitor the mutant actin isoform, TS-actin was mutated and transformed into a Gl14-3-3-HA expressing parasite line. In Gl14-3-3-HA immunoprecipitation, both TS-actin^R62D^ and control TS-actin co-precipitated in similar ratios as compared with endogenous GlActin (Figure 3F, 3G). This result suggests that the association between GlActin and Gl14-3-3 promoted by calyculin depends on increased actin phosphorylation rather than an increased amount of monomeric actin. To confirm that 14-3-3 complexes with phosphorylated actin, pulldowns of 14-3-3-TS were performed and the phosphorylation state of the associated actin was assessed with Phos-tag gels and western blotting. Indeed, compared with input, the phosphorylated forms of actin were enriched in 14-3-3-TS pulldown (Figure 3H); however, most of the co-immunoprecipitated actin was not phosphorylated. The presence of both phosphorylated and unphosphorylated actin in association with 14-3-3 indicates multiple modes of 14-3-3 association, which could include both direct binding and recruitment through actin binding proteins.

Since modulating actin phosphorylation changed the balance between F and G-actin, we anticipated that this would be apparent as changes in cellular actin organization. To verify this hypothesis, cells were treated with calyculin A or staurosporine for 30 minutes and then stained for GlActin and Gl14-3-3-HA or tubulin. Treatment with the phosphatase inhibitor calyculin A resulted in an apparent decrease in the robustness of actin structures and enrichment of Gl14-3-3 along the intracytoplasmic axonemes of the anterior flagella (Figure 4A). More severely impacted cells (27%, 83/300) lost cytoskeletal organization and became spherical (Figure 4B). Conversely, treatment with staurosporine led to increased cortical actin and brightly labeled F-actin structures, the most apparent of which are at the anterior of the cell (Figure 4A). Prominent actin filaments associated with the nuclei of staurosporine-treated cells were also apparent (Figure 4B and C). In 30% (90/300) of staurosporine treated cells, we observed an aberrant structure containing actin and 14-3-3 in proximity to the ventral disc, suggesting that membrane trafficking is also impaired (Figure 4C, asterisk). Nuclear size was also increased. Staurosporine treatment led to prominent nuclear actin filaments and a corresponding 89% increase in nuclear area compared to DMSO-treated control cells (p<0.001, n=43 Ctrl, n=32 Staur). This phenotype suggests that actin regulates nuclear size in *Giardia*. Overall, phosphorylation of the cytoskeleton appears to be correlated with Gl14-3-3 association and cytoskeletal disassembly while inhibiting phosphorylation with staurosporine stabilized cytoskeletal structures.

**Figure 4.**
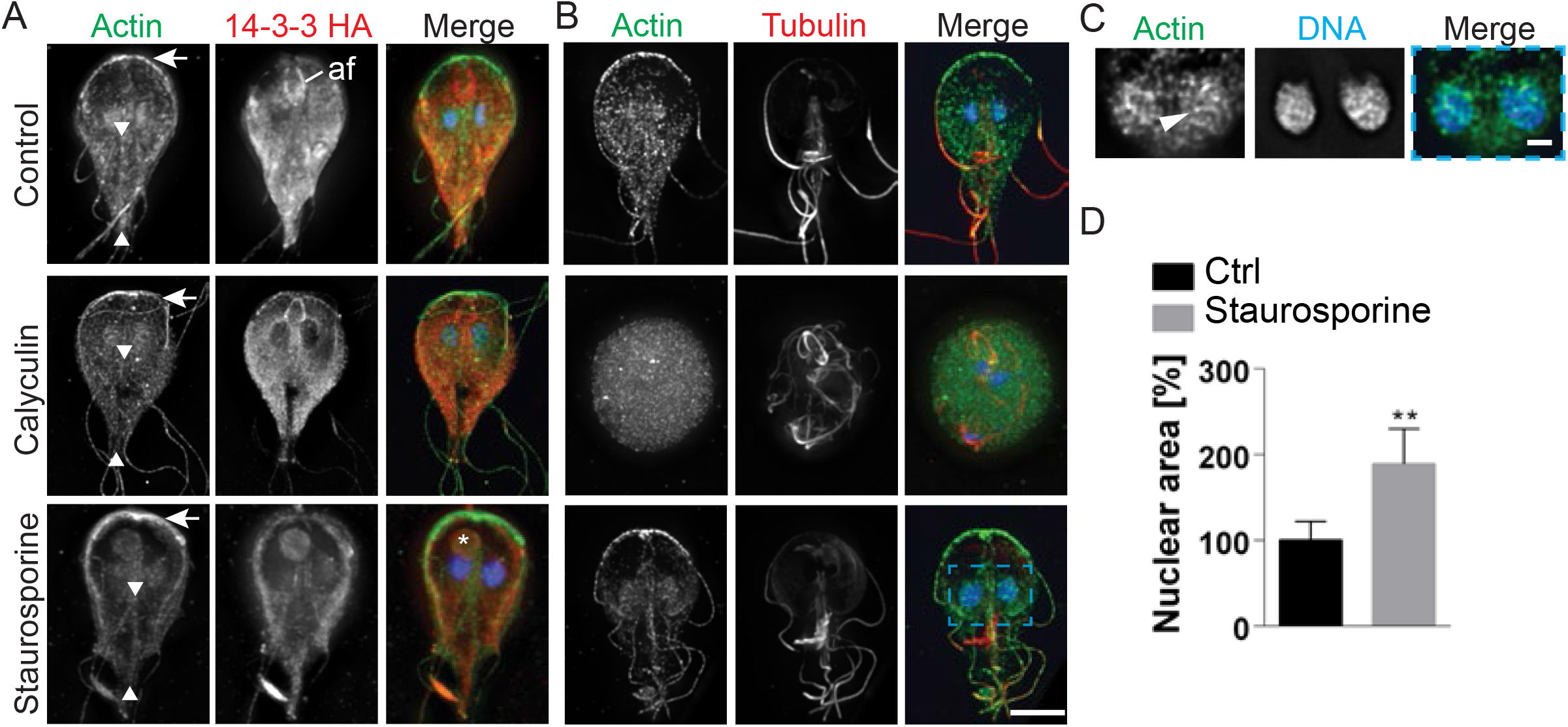
Filamentous actin structures are depleted by calyculin A treatment and enhanced by staurosporine treatment. (A) Projected images of actin (green), Gl14-3-3-HA (red), and DNA (blue) in the presence of calyculin A and staurosporine. Arrows marks the anterior of the cell where actin intensity is reduced by calyculin A treatment (increased phosphorylation) but enhanced by staurosporine treatment (reduced phosphorylation). Arrowheads mark the intracytoplasmic caudal flagella axonemes which are typically associated with actin, note calyculin A treatment resulted in loss of actin association with this structure while staurosporine treatment increased actin association. Calyculin A treatment enriched Gl14-3-3-HA along the intracytoplasmic axoneme of the anterior flagella (af). Asterisk marks the aberrant structure found in 30% of staurosporine treated cells. Note this structure is associated with the bare region of the disc, a conduit of cellular trafficking. (B) Projected images of actin (green), tubulin (red), and DNA (blue) in the presence of calyculin A and staurosporine. Note that 27% of calyculin A treated cells lost cytoskeletal organization and adopted a rounded cell shape. Scale bar= 5 μm. Nuclear area increased after staurosporine treatment. (C) A single optical section enlarged from blue box in B showing actin filaments associated with the nuclei, arrowhead marks a prominent filament. (D) Nuclear area quantified after treatment with staurosporine (mean ± SD, **p<0.01). Scale bar= 1 μm.

Next we reasoned that if Gl14-3-3 acts as a negative regulator of actin filament formation through sequestration of actin and/or actin regulatory proteins in *Giardia*, Rho GTPase signaling should act upstream of 14-3-3-actin complex formation. Expression of an inducible N-terminally HA-tagged constitutively active Q74L GlRac (HA-Rac^CA^; equivalent to Q61L Rac1) was previously observed to increase overall actin fluorescence, which we suggested was due to increased filament formation (21). Indeed, detailed analysis revealed the formation of prominent GlActin filaments at the cell cortex (Figure 5A). Induction of the HA-Rac^CA^ mutant protein lead to a decrease in the amount of actin that co-immunoprecipitated with 14-3-3-VSVG (Figure 5B). This is consistent with 14-3-3 acting as a negative regulator of filament formation where the monomeric 14-3-3 actin complex(s) must be reduced to facilitate filament formation. This result provides the first insight toward understanding how Rho GTPase signaling regulates actin in *Giardia* without any of the conserved proteins that normally link Rho GTPases and the actin cytoskeleton.

**Figure 5.**
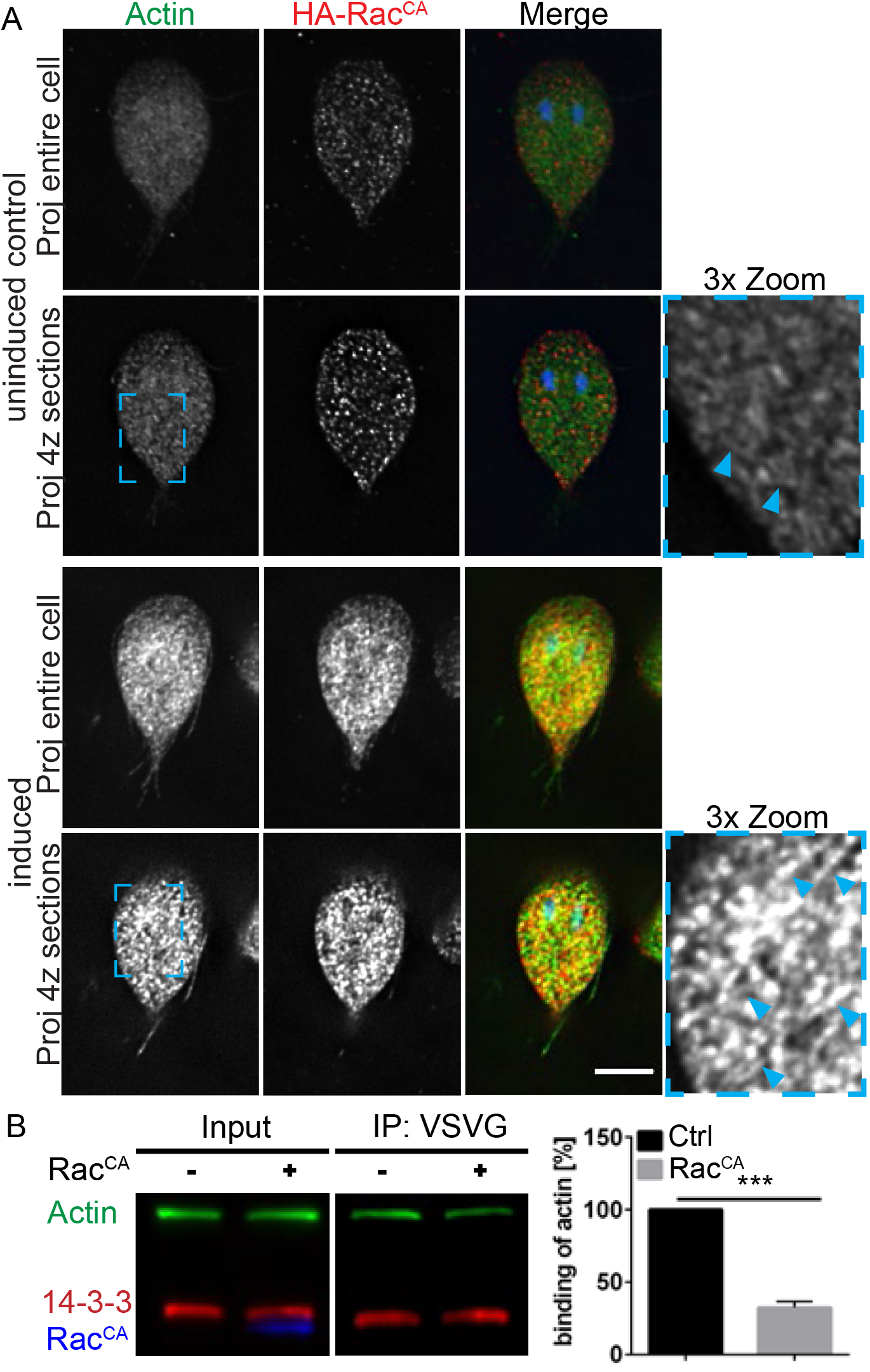
Rac signaling modulates 14-3-3-Actin complex formation. (A) Actin filaments (green) are more prominent in Tet-induced HA-Rac^CA^ expressing cells. Note that the *tet* promoter is leaky and some expression is detected in uninduced control cells (images scaled equally). (B) Immunoprecipitation of actin with 14-3-3-VSVG from uninduced (Rac^CA^ -) and induced (Rac^CA^ +) HA-Rac^CA^ cell lines and quantification of actin binding from three independent experiments (P<0.001). Scale bar= 5 μm.

After demonstrating that 14-3-3-actin complex levels could be modulated, we sought to determine if there are binding motifs in GlActin that could support direct interaction with Gl14-3-3. The 14-3-3-Pred prediction algorithm tool identified S330 (RVRIpSSP) and S338 (RKYpSAW) as the highest scoring predicted interaction sites (35). In agreement, the same sites were previously reported using a custom algorithm to identify binding sites in putative *Giardia* 14-3-3 interactors (22). These sites are similar to the canonical mode 1 site RXXp(S/T)XP [where p(S/T) are phosphorylated serine or threonine residues]; while not a perfect match, many 14-3-3 interacting proteins have been found that lack canonical mode 1-3 binding motifs (3, 35, 36). To assess the potential involvement of these sites, TS-actin was mutated to generate a S330A/S338A double mutant. The wild type and mutant TS-Actin constructs were introduced into the endogenously tagged Gl14-3-3-HA parasite line. Phos-tag gel analysis of the S330A/S338A double mutant, confirms that at least one of these two sites is phosphorylated (Figure 6B, C). The double mutant as well as single point mutants had reduced capacity to coprecipitate 14-3-3-HA (Figure 6D). The mean reduction in 14-3-3 binding was similar for S330A, S338A, and S330A/S338A double mutant. The double mutant, however, showed more consistent reduction as noted by error bar size in Figure 6E. Incomplete disruption of the 14-3-3 interaction could indicate that part of the 14-3-3 recruitment is mediated by association with actin binding proteins, but could also indicate the presence of additional interaction sites. Thus we also tested the possible involvement of T162 (VTHpTVP), a conserved residue identified by Scansite 3 as the highest scoring 14-3-3 interaction site (37). However, mutation of T162 to alanine did not disrupt 14-3-3-actin interaction, consistent with structure homology modeling that suggested this residue was not surface accessible (Figure S3). These results are in line with S330 and S338 having a role in promoting 14-3-3 and monomeric actin complex formation. However, our inability to completely disrupt actin complex formation indicates additional means for 14-3-3 association with actin.

**Figure 6.**
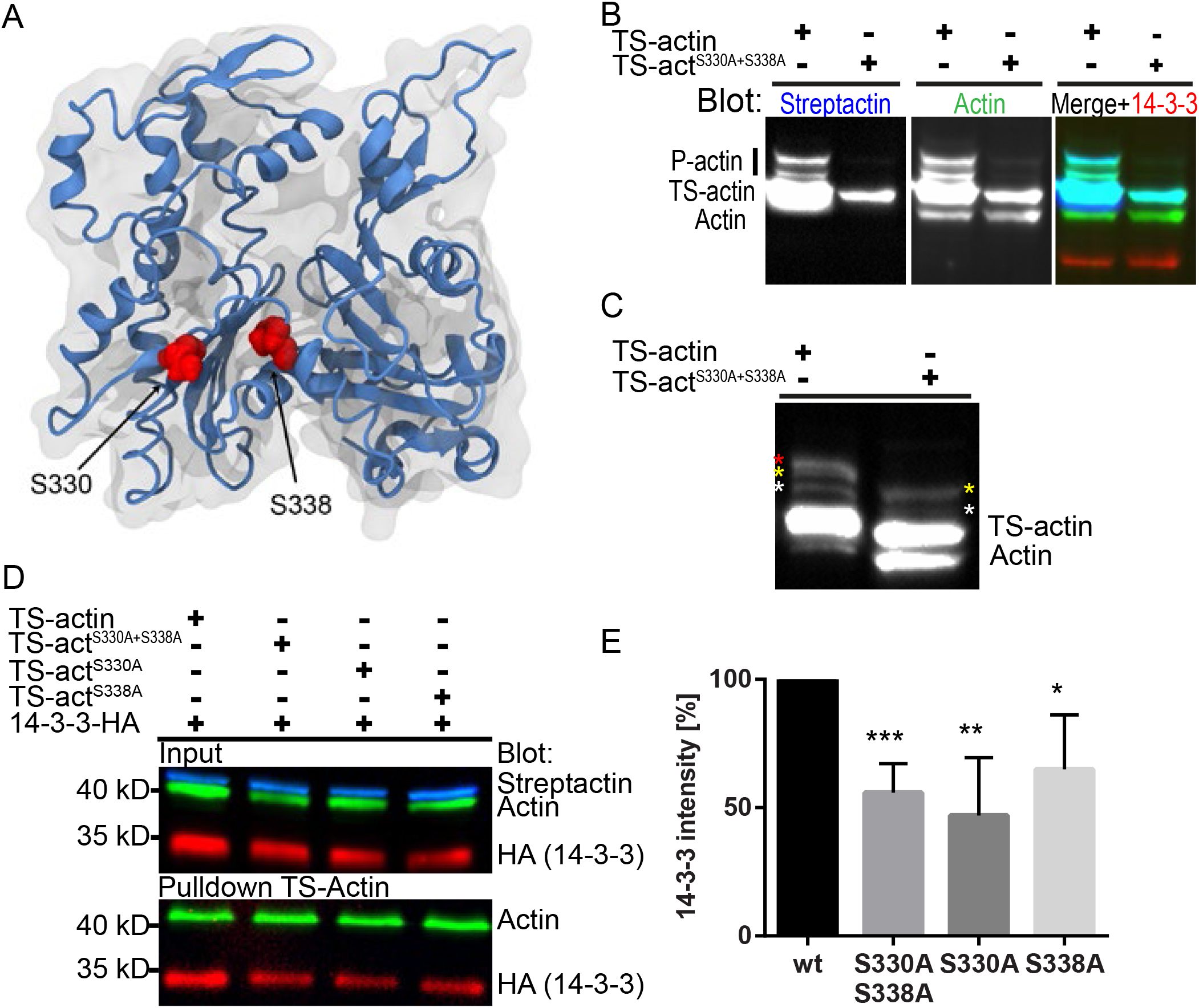
S330 and S338 of GlActin contribute to 14-3-3 complex formation. (A) Model of GlActin showing the position of S330 and S338 in an actin monomer. (B) Multiplexed immunoblot of total *Giardia* extracts after Phos-tag phosphate-affinity electrophoresis comparing phosphorylation of TS-actin and TS-actin^S330A+S338A^; anti-GlActin (green), StrepTactin-HRP (blue), and anti-HA (red). Note equal loading as indicated by 14-3-3 levels. (C) Samples from B overloaded and probed with anti-GlAactin antibody. (D) Affinity pulldown of TS-Actin variants blotted for GlActin and 14-3-3-HA. (E) Quantification of three independent affinity pulldown experiments show S330 and S338 contribute to 14-3-3 association (*p<0.05, * * p<0.01, * * * p<0.001).

To test whether Gl14-3-3 can directly bind GlActin, overlay experiments were performed using a 6XHis-GlActin expressed in and purified from *Giardia* extracts to ensure native phosphorylation. The overlay was performed with either the recombinant wild type GST-fused Gl14-3-3 or with the mutant GST-K53E, previously shown to be binding defective (38, 39). Binding of GST-Gl14-3-3 near the molecular weight of 6XHis-GlActin was not observed (Figure 7A). Instead, GST-Gl14-3-3 bound to four bands on the blot corresponding to proteins co-purified with 6XHis-GlActin. The most prominent labeling is associated with a major co-purified protein at 80 kD also seen in silver staining. Immunoblotting with an anti-pSer antibody confirmed that a fraction of actin was phosphorylated, as well as some co-purified proteins, including the 80 kD protein band (Figure 7A).This result confirms that some of the 14-3-3-actin interaction is due to recruitment through additional binding partners that can mediate the formation of a complex containing both Gl14-3-3 and GlActin. The inability to bind directly to actin could reflect a requirement for actin to be natively folded or that the interaction site is low affinity and additional proteins are needed to stabilize binding (40).

**Figure 7.**
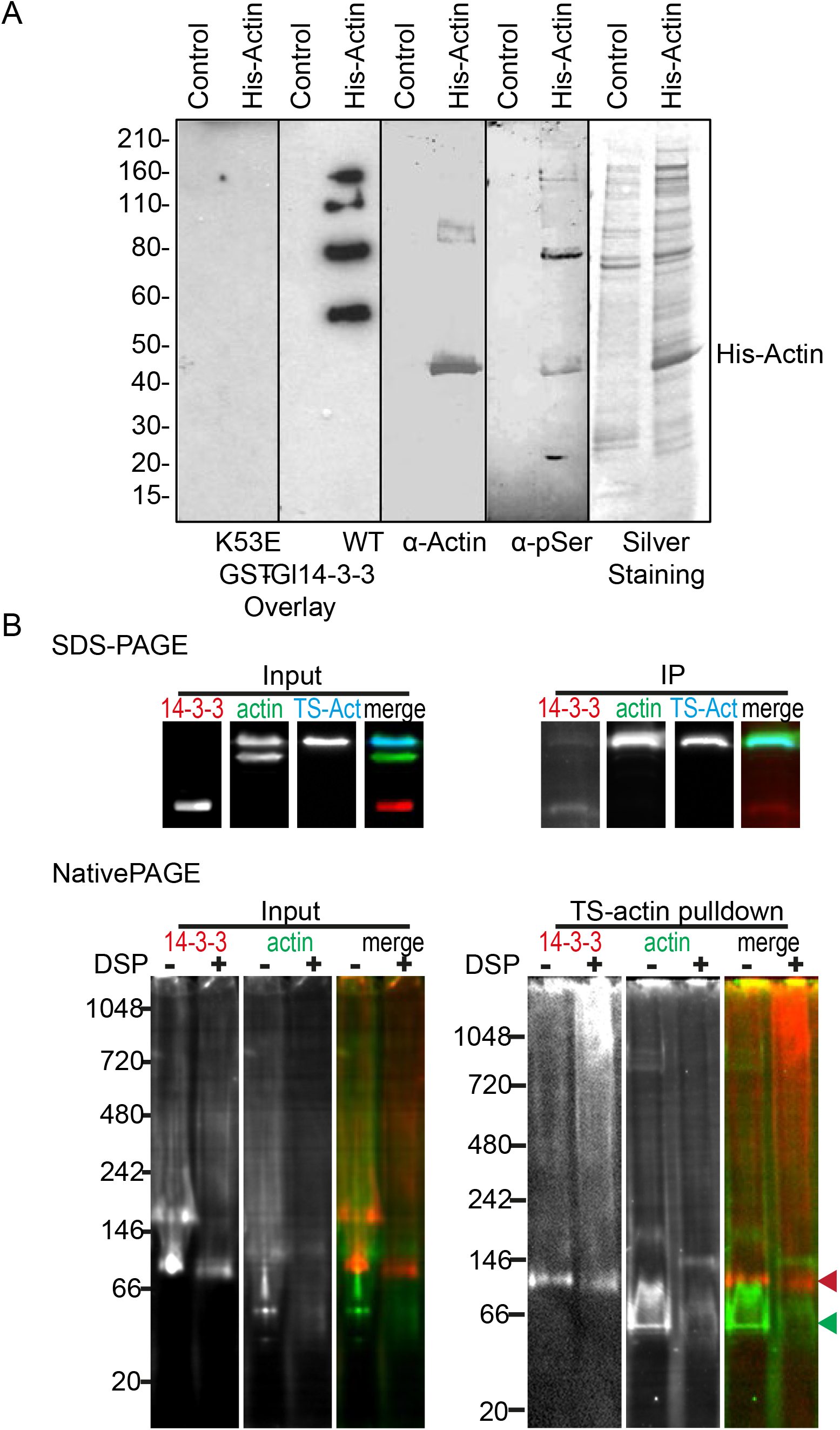
14-3-3-actin complex formation requires intermediate proteins. An immunoblot of affinity purified 6xHis-GlActin (His-Actin) or mock purification from wildtype trophozoites was assessed by overlay with recombinant GST-Gl14-3-3 or the 14-3-3 binding defective mutant GST-K53E. Interaction of GST-Gl14-3-3 with actin and co-purified proteins was revealed by incubation with anti-GST-HRP. The same membrane was stripped and probed with mouse anti-GlActin and again with mouse mAb anti-pSer. Silver stained protein purifications are shown in the last inset. Molecular size markers (kDa) are on the left. The position of His-Actin is indicated on the right. (B) Affinity purified TS-actin was run on SDS Page and NativePAGE. The native gel analysis includes DSP crosslinked samples to preserve native complexes. Note that DSP treatment reduces the amount of ~100kD dimeric 14-3-3 (red arrowhead) and increases the high molecular weight smear. The position of putative monomeric TS-actin (45.2 kD expected size) is marked with a green arrowhead.

To assess whether *Giardia* contains a 14-3-3 complex appropriately sized for direct interaction, we turned to native gel electrophoresis. If a 14-3-3-actin complex formed without any additional proteins, it would contain a dimer of 14-3-3 (41) and a single TS-actin monomer. Previous native gel analysis indicates that 14-3-3 dimers run at ~80 kD (41) and TS-actin is 45.2 kD; taking into account the size of the 3XHA tag (3.8KD) the anticipated complex size for direct binding is around 130 kD. TS-actin and associated proteins were purified and immediately run on native gels and transferred to membrane for western blotting. A prominent band of 14-3-3 was detected running between the 66 and 140 kD native gel markers (Figure 7B). However, a similarly sized band is also apparent in the input, suggesting this band represents free 14-3-3 dimer and that the complexes were unstable. In an effort to stabilize actin complexes, we used the membrane permeable crosslinker DSP to crosslink samples before lysis. Crosslinking did change the distribution of actin and a prominent actin band running just below 146 kD was enriched in the pulldown (Figure 7B); however, a corresponding enrichment of 14-3-3 was not apparent. Instead, the higher molecular weight smear of 14-3-3 became more prominent, consistent with multiple actin interactors having the capacity to bind actin. Future work will be required to identify these 14-3-3-actin interacting proteins and determine their specific roles in regulating the actin cytoskeleton. Although the relationship between 14-3-3 and actin appears to be more complicated than we initially imagined, our results indicate that 14-3-3 plays a decisive role in actin regulation.

## Discussion

Proteomic studies support complex formation between 14-3-3 and actin without the involvement of cofilin (10-12, 22). A recent study of 14-3-3σ function in the basal-like progression of breast cancer cells found that 14-3-3σ forms a complex containing actin and two intermediate filament proteins (13). Analogous to what we have reported here, the 14-3-3σ complex sequesters soluble actin in a bioavailable form that is then used for directional assembly of the cytoskeleton during cell migration. Similarly, *Giardia* appears to use Gl14-3-3 to restrict actin assembly at specific subcellular regions as evidenced by the 14-3-3 knockdown experiments where actin filaments were dispersed throughout the cell and typical actin organization was lost.

We estimated the intracellular concentration of actin to be around 4.7 μM. This relatively low concentration is nevertheless above the critical concentration for GlActin (21), indicating the need for actin sequestration. The emerging view of actin network homeostasis is that sequestering proteins play an important role in partitioning actin to competing F-actin networks (42, 43). In other eukaryotes, the concentration of sequestering proteins exceed the actin monomer pool (44), but *Giardia* lacks all known monomer sequestering proteins. Our results indicate that 14-3-3 is associated with monomeric actin in complexes with other interaction partners (Figure 7) and that complex formation has a reverse correlation with actin filament formation (Figure 3, 4, 5). Therefore, it seems likely that 14-3-3 functions in the partitioning of actin complexes for sub-functionalization in addition to having a role in maintenance of G/F-actin homeostasis.

Complex formation between 14-3-3 and actin appears to be both phosphodependent and independent. The putative 14-3-3 phospho-binding motif centered on S338 is conserved in mammalian actin and has previously been implicated as an AKT phosphorylation site (45). The 14-3-3 binding site prediction tool 14-3-3PRED identified S338 as the highest scoring site for both GlActin and human β-actin. We have shown this site as well as S330, contribute to 14-3-3 recruitment and at least one of these predicted interaction sites is phosphorylated as shown by a change in mobility on Phos-tag gels. Our ability to perform biochemical assays with *Giardia’s* highly divergent actin remains limited. Overlay assays and native gels did not provide support for direct interaction between 14-3-3 and actin. However, it remains possible that some of the observed complexes contain 14-3-3 directly bound to actin with additional interactors that work to stabilize the complex (40). Indeed the 14-3-3σ complex contains two intermediate filament proteins, yet *in vitro* actin polymerization assays containing only actin and 14-3-3 demonstrated that 14-3-3 could directly regulate actin dynamics (13). The need for additional proteins to stabilize the 14-3-3 interaction with S330 and S338 would reconcile our data; however, a caveat is that these point mutants could have disrupted actin folding and therefore disrupted the binding of proteins that recruit 14-3-3. While we have yet to pursue proteins that associate with both 14-3-3 and actin, independent proteomic studies aimed at identifying 14-3-3 and actin interactors point towards proteins of interest (19, 22). Besides actin and 14-3-3 there are 14 proteins in common between the two studies (Table S1). Eight of these proteins belong to the highly conserved Chaperonin containing TCP-1 (CCT; MW 56.3-64.7 kD) which is known to have a role in actin folding (46). The TCP-1 epsilon subunit has been implicated in the regulation of actin dynamics and this subunit has two, albeit untested, canonical mode 1 14-3-3 recognition motif (22, 47). Intriguingly, the epsilon subunit was the most abundant component of the TCP-1 complex found in our actin interactome. The list also includes TIP49 (GL50803_9825, 51.4 kD), which we have validated as a robust actin interactor through reciprocal pulldown (19). Other proteins include an SMC domain protein (GL50803_6886, 102.8 kD), two dynein heavy chain proteins (GL50803_111950, 570.3 kD; GL50803_42285, 834.7 kD), and two proteins without any conserved domains (GL50803_15251, 32.5kD; GL50803_5120, 26.6 kD). All of these proteins have at least one canonical mode 1 14-3-3 recognition motif but whether they are involved in recruiting 14-3-3 to actin complexes remains to be determined.

Overall, our results support a role for 14-3-3 associating with monomeric actin complexes that are downstream of GlRac, where phosphorylated actin and phosphorylated actin interactors are held inactive. This work illustrates the conserved role of 14-3-3 as an actin regulator albeit through an alternative set of actin binding proteins which remain to be identified and validated.

## Materials and Methods

### Strain and culture conditions

*Giardia intestinalis*, strain WBC6 was cultured as in (48).

### Culture and Microscopy

Fixations were performed as in (21). Anti-HA HA7 (Sigma), anti-tubulin 11-B6-1 (Sigma), and anti-GlActin 28PB+1 (21) antibodies were used at 1:125, secondary antibodies were used at 1:200 (Molecular Probes). Isotype specific anti-mouse secondary antibodies (Molecular Probes) were used for co-localization of tubulin (IgG2b) and HA (IgG1). Images were acquired on a DeltaVision Elite microscope using a 100x 1.4 NA objective and a Coolsnap HQ2 or PCO EDGE sCMOS camera. Deconvolution was performed with SoftWorx (GE, Issaquah, WA). Average and maximal projections were made with ImageJ (49) and figures were assembled using Adobe Creative Suite (Mountain, CA). Nuclear area measurements were made from thresholded maximal projections of DAPI staining using ImageJ.

### Morpholino and Drug Studies. Morpholino and Drug Studies

Morpholino treatments with anti-14-3-3 CGCGTAAATGCCTCGGCCATAGGTT and control CCTCTTACCTCAGTTACAATTTATA were performed as in (50). Calyculin A and Staurosporine (LC Laboratories, Woburn, MA) were diluted in DMSO and used at 1 μM and 200 nM final concentrations, respectively. Cells were treated for 45 min at 37^o^C.

### Constructs and mutagenesis

Endogenous tagging of Gl14-3-3 with 3XHA and construction of N-terminally TS tagged GlActin are described in (19). Actin T162A, S330A and S338A site mutations were introduced into TwinStrep-Actin using QuikChange Lightning Multi Site-Directed Mutagenesis Kit (Agilent Technologies, Wilmington, DE) using primer S330: ctgtcctcgggactagctatgcgcacacgcttc, T162: gacggggtgacgcatgctgttcctgtgtac, and S338: cgaggacagaaagtacgctgcctgggttggtg. Construction of Q74L HA-GlRac (HA-Rac^CA^) under *tet* promoter is described in (21). To construct 14-3-3-VSVG, 14-3-3 (GL50803_6430) was cut from pKS-6430-3HA_Neo vector (19) using *BamHI* and *Afl II* and cloned into pKS-VSVG_Neo. pKS-VSVG_Neo was constructed as follows. VSVG tag flanked with 5′ *Afl II* and *EcoR I* 3′ restriction sites was PCR amplified using VSVG F′ 5′ TGGCTTAAGTATACTGATATTGAAATGAATCGCTTAGGTAAAGGGTCCTACACCGACATCGAGATGAACCGCTTG 3′ and VSVG STOP R′ 5′ TTTGAATTCTCATTTTCCAAGTCTGTTCATTTCTATGTCTGTATAAGAGCCCTTGCCCAAGCGGTTCATCTC GATGTC 3′ overlapping template oligos and cloned into pKS-3HA.neo to replace the 3xHA tag. A multiple cloning site was introduced into the vector by ligation with annealed linker (MCSlinkerF: 5′ gatcccccgggctgcaggaattcgatatcaagcttatcgataccgtcgacctcgagc 3′; MCSlinkerR: 5′ ttaagctcgaggtcgacggtatcgataagcttgatatcgaattcctgcagcccgggg 3′). To ensure integration into the genome and endogenous levels of expression, the 14-3-3-VSVG NEO vector was linearized using *BsRGI*, EtOH precipitated and transformed into *Giardia* cell line already containing Q74L HA-GlRac under the *tet* promoter. To construct 14-3-3-TS, 14-3-3 was amplified from genomic DNA using Forward OL368 5′ tatagaatactcaagcttggcgcgccGGAAAATGTGTGATCACCCC 3′ Reverse OL369 5’ GCTCCAAGCGCTCCCaccggtCTTCTCCTCGGCATTATCGT 3′ primers and the product was inserted into p7031 TwinStrepPac digested with AgeI and AscI using Gibson assembly. The 6XHIS-GlActin vector for expression in *Giardia* was generated by PCR amplifying the 6XHis-tag and coding region from 6XHis-Actin Bestbac (21) using Primers 408176xHis-F NheI atggctagccatcaccatcaccatcacga and 408176xHis-R ClaI tgtatcgataacaatcccgg. The PCR amplified insert was ligated into the TwinStrep-Actin vector (19) after preparing the vector and PCR product with *Cla I* and *Nhe I*.

### Detergent extractable actin

This assay compares the fraction of Triton X-100 extractable actin versus non-extractable actin within the cell. While some filaments may be extracted by detergent treatment, the detergent-extractable fraction is largely composed of monomeric actin while actin filaments associated with larger scaffolds remain within the cell (51, 52). Confluent 8 ml cultures were treated for 45 minutes with drugs, chilled to detach cells, and then pelleted at 700xg. The pellet was re-suspended in 800 μl of HBS plus protease inhibitors and moved to a microfuge tube. The cells were pelleted again and then resuspended in 100 μl of actin stabilizing lysis buffer 50 mM PIPES (pH 6.9), 50 mM NaCl, 5 mM MgCl_2_, 5 mM EGTA, 5% (vol/vol) glycerol, 0.1% Triton X-100, 0.1% Tween 20, 0.1% 2-mercapto-ethanol, 0.2 mM ATP, and HALT protease inhibitor. The samples were incubated on ice for 5 minutes to perforate the membrane and then pelleted at 700xg for 5 minutes. The supernatants were reserved and the pellets were re-suspended in 80 μl of 8 M urea for one hour on ice. Equal amounts of each sample were boiled in sample buffer, loaded on SDS PAGE, and actin levels were analyzed by immunoblotting.

### Immunoprecipitation

Immunoprecipitation began with 1-3 confluent 13 ml tube per cell line. After detachment by icing, cells were pelleted at 700xg and washed once in HBS. The cells were resuspended in 300 μl of lysis buffer (50 mM Tris (pH 7.5), 150 mM NaCl, 7.5% glycerol, 0.25 mM CaCl_2_, 0.25 mM ATP, 0.05 mM DTT, 0.5 mM PMSF, 0.1% Triton X-100, 2X Halt Protease inhibitors (Pierce)) and sonicated. The lysate was cleared by centrifuging at 10,000xg for 10 minutes at 4°C and added to 30 μl of lysis buffer-equilibrated Anti-HA (Sigma), anti-VSVG (Sigma) or StrepTactin (IBA) beads. After 1.5 hours of binding the beads were washed four times with wash buffer (25 mM Tris (pH 7.5), 150 mM NaCl, 0.25 mM CaCl_2_, 0.25 mM ATP, 5% Glycerol, 0.05% Tween-20) and then boiled in 50 μl of sample buffer.

### Phos-Tag Gels

After lysis or immunoprecipitation, samples were loaded onto 10% SDS gel supplemented with 100 μM MnCl2 and 20 μM acrylamide-pendant Phos-tag^TM^ AAL-107 (NARD Institute, Ltd.; from 5 mM stock solution in 3% MeOH in distilled water, prepared according to manufacturer′s instruction) to detect mobility shift of phosphorylated proteins. The gels were run at 100 V for approximately 2 hours.

### Native Gels

TS-Actin and associated proteins were purified as above with slight modifications. After cell lysis the complexes were bound to StrepTactin resin (IBA Lifesciences, Germany). Unbound complexes were washed out two times and the 14-3-3-actin complexes were eluted with wash buffer supplemented with 2 mM biotin (Sigma). The samples were dissolved in NativePAGE™ Sample Buffer (4X) and immediately loaded onto 4-16% NativePAGE™ Novex^®^ Bis-Tris Mini gel (Life Technologies) according to the manufacturer’s instructions.

### Western Blotting

After gel electrophoresis, proteins were transferred to Immobilon-FL using wet transfer at 200 mA for 1 hour (2 hours for Phos-Tag and native gels) and blocked in TBS+5% nonfat dry milk. IPs with the double transformant 14-3-3-VSVG and tet inducible HA-Rac^CA^ cell line were performed after 24 hours of 20 ug/ml tetracycline induction. Primary antibodies 28PB+1 (actin, rabbit polyclonal) was used at 1:2500, Sigma HA7 (HA, IgG1) was used at 1:2500, IBA StrepTactin-HRP used at 1:7 000, Sigma P5D4 (VSVG, IgG1) used at 1:1 500, Abcam HA.C5(IgG3) was used at 1:1500 and detected with Molecular Probes fluorescent isotype specific secondary antibodies at 1:2500. For quantification of cellular Gl14-3-3 concentration parasites were lysed in PBS/1% Tritox X-100 (supplemented with protease and phosphatase inhibitors) for 1h on ice, pelleted 15 min at 13,000 rpm at 4°C, and supernatant collected and quantified by Bradford assay. Recombinant GST-Gl14-3-3polyG20 was prepared and purified as previously described (41). WB was performed with anti N-terminal Gl14-3-3 rabbit antiserum (39) 1:10 000. For quantification of cellular GlActin concentration, trophozoites were washed 2x with HBS + HALT protease inhibitor and PMSF, then boiled in 2% SDS 62.5 mM TrisHCl pH 6.8. Protein concentration in lysates were then quantified by DC Assay. TwinStrep-tagged actin was purified from *Giardia* trophozoites. Trophozoites were lysed by sonication in the following buffer: 50 mM TrisHCl pH 7.5, 150 mM NaCl, 7.5% glycerol, 0.25 mM CaCl2, 0.25 mM ATP, 0.1% Triton with PMSF and HALT protease inhibitors. Lysates were rotated with StrepTactin resin (IBA) for 2 hours at 4 °C. Resin was washed a total of four times with modified G-Buffer (10 mM Tris 8.0, 0.2 mM CaCl_2_, 0.2 mM ATP). Washes 1, 3, and 4 consisted of G-Buffer with 500 mM NaCl and 0.1 mM DTT. Wash 2 consisted of G-Buffer plus 500 mM NaCl, 0.1 mM DTT, and 0.1% Tween 20. Protein was eluted with 2.5 mM biotin in G-Buffer. Western blot was performed with polyclonal GlActin rabbit antibody (21).

### Overlay assays

*Giardia* protein extracts were prepared from 1×10^9^ WBC6 trophozoites or 6XHIS-GlActin transgenic line. Parasites were recovered by chilling on ice, then washed three times with cold PBS and the cell pellet was frozen at -70 °C overnight. Cells were resuspended in 1 ml of buffer A (50 mM NaH_2_PO_4_, 300 mM NaCl, 10 mM imidazole, 0.005% Tween 20, pH 8.0), supplemented with protease/phosphatase inhibitor cocktail (Cell Signaling, Danvers, MA, USA), lysed by 7 cycle of sonication at 15% of power (Sonoplus, Bandelin) and, centrifuged at 24.000 rpm at 4°C for 30 min. Supernatant was collected and incubated with 200 μl of Ni-NTA Magnetic Agarose bead suspension (Qiagen, Germany) at 4°C for 1 h with gentle rotation. Beads were extensively washed with wash buffer A containing 20 mM imidazole. Bound proteins were eluted in buffer A supplemented with 250 mM imidazole and then dialyzed and concentrated in G buffer (50 mM Tris 8.0, 2 mM CaCl_2_, 2 mM ATP). An aliquot (1:4) of purified proteins was separated on 4-12% NuPAGE gel in MOPS-SDS buffer (Invitrogen), blotted on nitrocellulose membrane and blocked 1h in 5% nonfat dry milk/HT buffer (20 mM Hepes–KOH, pH 7.6, 75 mM KCl, 5 mM MgCl_2_, 1 mM DTT, 0.1 mM EDTA, 0.04% Tween 20). The membrane was incubated with 10 μg/ml of either GST-Gl14-3-3 or the GST-K53E mutant (Lalle et al., 2006; Lalle et al., 2010) in 2.5% nonfat dry milk/HT buffer over night at 4°C. Protein-protein interaction was assessed by incubation with anti-GST-HRP (GE Healthcare, 1:1000) and revealed by chemiluminescence. The membrane was then stripped and re-probed with anti-GlActin (1:5000) and western blot developed using DAB. Alternatively, the membrane was probed with mouse mAb anti-pSer (Sigma-Aldrich, 1:200) in 3% BSA/TTBS buffer and western blot developed using DAB. Aliquots (1:20) of purified protein were alternatively visualized by silver staining (GE Healthcare).

### Statistical Analysis

For each experiment with a p-value, we compared at least three biological replicates using a two-tailed t-test.

### Structure Analysis

We used the Oda et al. F-actin structure (pdb 2ZWH (53)) and the ADP G-actin structure (pdb 1J6Z (54)) to determine the solvent accessibility of T162, S330 and S338.

## Acknowledgments

We thank B. Wakimoto, T. Yamaki and members of the Paredez lab for critical reading of this manuscript. This work was supported by NIH 1R01AI110708-01A1 to A.P. and NPUI (LO1417) of the Czech Ministry of Education, Youth and Sports to J.K.

